# Snake venom gland cDNA sequencing using the Oxford Nanopore MinION portable DNA sequencer

**DOI:** 10.1101/025148

**Authors:** Adam D Hargreaves, John F Mulley

## Abstract

Portable DNA sequencers such as the Oxford Nanopore MinION device have the potential to be truly disruptive technologies, facilitating new approaches and analyses and, in some cases, taking sequencing out of the lab and into the field. However, the capabilities of these technologies are still being revealed. Here we show that single-molecule cDNA sequencing using the MinION accurately characterises venom toxin-encoding genes in the painted saw-scaled viper, Echis coloratus. We find the raw sequencing error rate to be around 12%, improved to 0–2% with hybrid error correction and 3% with de novo error correction. Our corrected data provides full coding sequences and 5′ and 3′ UTRs for 29 of 33 candidate venom toxins detected, far superior to Illumina data (13/40 complete) and Sanger-based ESTs (15/29). We suggest that, should the current pace of improvement continue, the MinION will become the default approach for cDNA sequencing in a variety of species.

## Background

The transcriptome can be defined as all of the RNA molecules expressed by a cell or population of cells, for example in a particular tissue [1]. As this includes all expressed mRNA molecules, the transcriptome can be inferred to represent all protein coding genes that are actively transcribed at the time of sampling [2]. In theory then, the transcriptome is the precursor to the proteome of a cell or tissue, although post-transcriptional and post-translational modification and regulation are likely to cause some disparity between the two. Traditionally transcriptomes were analysed via cloning and sequencing of expressed sequence tags (ESTs) whereby short fragments of a cDNA library are sequenced and clustered to give a contiguous sequence. ESTs are ultimately limited by their short length (typically 200-800bp) [3] and low coverage, meaning lowly expressed transcripts and splice variants are likely to remain undetected [2]. The advent of “next-generation” sequencing technologies such as the Roche 454, ABI SOLiD and Illumina Genome Analyzer platforms in the first decade of the 21^st^ century facilitated a step-change in transcriptome studies: increased sequencing depth improves the likelihood of recovering full-length transcript sequences (including lowly expressed transcripts), and higher resolution aids in the identification of splice variants. As the number of reads sequenced from a particular transcript will be representative of the amount of that transcript present in a sample, such data is also quantitative [4]. Both the ABI SOLiD and Roche 454 systems are no longer available/supported, and the DNA sequencing market is now largely dominated by platforms that produce high numbers of short reads. The assembly of these reads into full transcript sequences poses several challenges, especially in the absence of a reference genome. Unlike the genome (which remains relatively static), the transcriptome can be highly variable, with mRNA transcripts encoding different genes present at different abundances within a given sample, resulting in uneven sequencing coverage [2, 5], particularly in highly transcriptionally active tissues. The short read length also means that reads from highly similar transcripts, such as paralogs (members of a gene family produced by gene duplication, as distinct from orthologs which are produced via speciation) belonging to the same gene family, may be fused during the assembly process resulting in chimeric sequences. Alternative transcripts of the same gene may be omitted altogether if the abundance of one variant in a sample significantly outweighs the other(s) [6] and, finally, shared homologous sequences in related genes may be incorporated or omitted erroneously, especially if they are highly conserved.

The characterisation of the venom gland transcriptomes of venomous snakes has been particularly useful in revealing the genetic basis of inter-and intra-specific variation in venom composition, something which has significant implications for antivenom manufacture [7–10]. Although genome sequences for some venomous species are now available (including the king cobra *Ophiophagus hannah* [11] and the speckled rattlesnake, *Crotalus mitchellii* [12]), for the vast majority of species *de novo* assembly of short-read sequences has been the only feasible (and cost-effective) approach. However, such approaches have difficulty in accurately reconstructing full-length sequences for highly similar paralogs in some key venom gene families. For example, we have previously found that assemblies of Illumina HiSeq data using Trinity (version trinityrnaseq_r2012-04-27, [13]) only provided full-length coding sequences for 13 candidate venom toxin encoding genes in the painted saw-scaled viper (*Echis coloratus*) ([14, 15]). Others have shown similar issues with venom gland transcriptomes from the Okinawa habu (*Protobothrops flavoviridis*) and the Hime habu (*Ovophis okinavensis*), where 37/103 and 29/95 complete transcripts were identified respectively [16]. Attempts have been made to develop an assembler specifically for samples containing large numbers of highly similar transcripts, such as VTbuilder [17], although the current version has an upper limit of 5 million ≥120bp reads, making it less suitable for the analysis of large-scale data generated from the most recent Illumina platforms or for the re-analysis of older datasets with shorter read lengths. Long-read data derived from single-molecule sequencing should eliminate many of the current problems associated with the investigation of snake venom gland transcriptomes, but the only currently commercially-available long-read platform (the Pacific Biosciences RSII) typically requires a large number of flowcells (12–16 for a comprehensive survey of full-length isoforms, each costing £400) and several size-selection and PCR steps. The Oxford Nanopore MinION (Figure 1) is a portable, USB 3.0-powered DNA sensing device that uses an application-specific integrated circuit (ASIC) to detect miniscule voltage changes resulting from the movement of DNA strands through pores embedded in a membrane. The disposable flowcell (£300-500 each depending on quantity purchased) contains 2,048 sensor wells (each of which contains a single pore), with 512 measurement channels below these. The choice of which is the “best” pore to use is performed by the multiplexer (or “mux”) during an initial platform QC step, and the standard 48 hour run protocol performs one switch to an alternative pore after 24 hours. A “motor” protein unwinds the DNA as it enters the pore and controls the speed at which the DNA translocates the pore to facilitate accurate base-calling and a “hairpin” adaptor at the other end of the DNA enables both strands to be read. Since the same piece of DNA is analysed twice, a consensus (“2D”) read of greater accuracy can therefore be generated. The MinION was initially made available to selected users in the MinION Access Program (MAP) in spring 2014, with the first publications emerging in late 2014/early 2015 and the rapid dissemination of results and protocols facilitated by an active online community and preprint servers such as bioRxiv (http://biorxiv.org). The utility of the MinION for the rapid and accurate investigation of disease outbreaks [18]; microbial diversity analysis [19]; sequencing of bacterial and viral genomes [19–22], haplotype resolution [23] and even for the characterisation of more complex eukaryotic genomes [24] has already been demonstrated. However, the utility of this device for the characterisation of transcriptomes has not yet been comprehensively investigated (a previous study investigating the venom gland transcriptome of the Okinawa habu (*Protobothrops flavoviridis*) was based on an amplicon sequencing protocol, and produced very small amounts of data from a single flow cell, (Table 1) [25]). We therefore set out to establish the feasibility of using the Oxford Nanopore MinION to characterise snake venom gland transcriptomes, something for which long-read data derived from single DNA molecules should be eminently suitable, and which should help to overcome the issues associated with *de novo* assembly of highly similar venom gene paralogs. We chose to investigate the painted saw-scaled viper, *Echis coloratus* (Figure 1), as this species is not only a member of the genus of snakes thought to be responsible for more deaths than any other [26, 27], but it is also one for which we have Illumina HiSeq data [14, 15] and for which ESTs derived from Sanger (dideoxy, chain-termination) sequencing are available [26].

**Figure 1.**
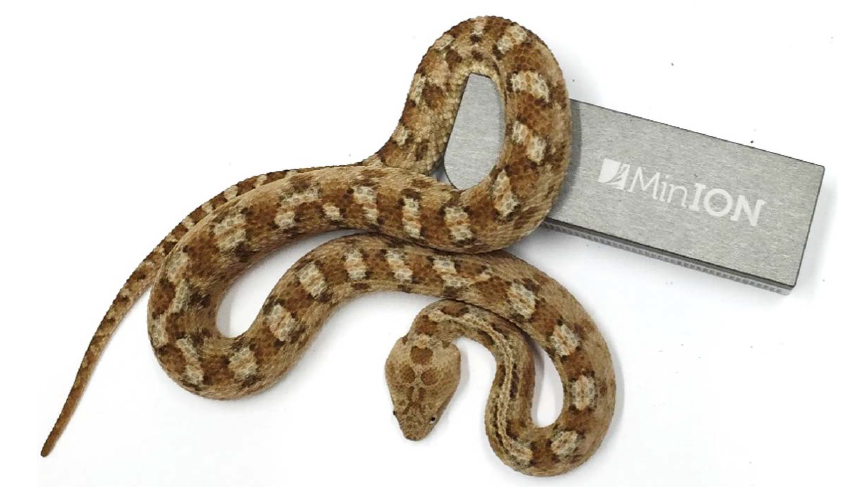
The Oxford Nanopore MinION portable DNA sequencing device and a painted saw-scaled viper, *Echis coloratus*.

**Table 1.** Oxford Nanopore MinION venom gland transcriptome sequencing statistics. Painted saw-scaled viper (*Echis coloratus*) data was derived from two individuals (Eco6 and Eco8), using four R7.3 flowcells and both the standard 48 hour run (with a “re-mux” voltage change at 24hrs) and a modified run utilising four re-mux steps at 8 hour intervals. *Protobothrops flavoviridis* statistics are derived from a reanalysis of the raw data of Mikheyev and Tin [25]. ‘Pass’ data is that selected by the base-calling software Metrichor as being high quality and consists entirely of 2D read data.

|  |  | Eco6<br>(48hr) | Eco8<br>(48hr) | Eco6<br>(4x 8hr) | Eco8<br>(4x 8hr) | *Protobothrops*<br>*flavoviridis* |
| --- | --- | --- | --- | --- | --- | --- |
|  | Available pores | 436 | 332 | 345 | 387 | (unknown) |
| All data | Total reads | 93,697 | 47,068 | 66,916 | 58,628 | 2,057 |
|  | Total bases (Mb) | 132.1 | 70.3 | 81.7 | 80.1 | 1.3 |
|  | Max length (bp) | 454,436 | 278,051 | 363,606 | 212,026 | 29,363 |
|  | Min length (bp) | 5 | 5 | 5 | 7 | 5 |
|  | Mean length (bp) | 1,410 | 1,493 | 1,220 | 1,378 | 614 |
|  | N50 (bp) | 1,577 | 1,753 | 1,412 | 1,648 | 823 |
| 'Pass' data<br>only | Total reads | 16,804 | 7,190 | 9,172 | 7,786 | 16 |
|  | Total bases (Mb) | 22.7 | 11 | 12.2 | 11.7 | 0.019 |
|  | Max length (bp) | 12,639 | 5,869 | 10,422 | 8,521 | 2,195 |
|  | Min length (bp) | 247 | 251 | 287 | 248 | 650 |
|  | Mean length (bp) | 1,352 | 1,536 | 1,333 | 1,509 | 1,220 |
|  | N50 (bp) | 1,536 | 1,801 | 1,504 | 1,782 | 1,323 |

## Results and Discussion

We used four R7.3 flowcells to characterise the venom gland transcriptome of *Echis coloratus*, using venom gland tissue samples from two individuals (“Eco6” and “Eco8”) for which we had previously generated data on the Illumina HiSeq platform [14, 15]. We used both the standard Oxford Nanopore 48 hour run script (which performs a voltage “re-mux” after 24 hours) and a set of modified scripts (John Tyson, pers com) which perform four re-mux steps at 8 hour intervals. Of the 512 theoretically available pores per flowcell, initial platform QC showed between 332-436 as actually being available for sequencing (Table 1) - figures within the range seen by many other participants of the MAP. Base-calling of data derived from the MinION is performed by cloud-based software called Metrichor and the resulting sequence data (in .fast5 format) is divided into ‘pass’ and ‘fail’ folders. The contents of the ‘fail’ folder are typically 1D and low-quality 2D data and the ‘pass’ folder contains only high-quality 2D reads. We have chosen to focus only on these high-quality ‘pass’ reads for our analyses. Our four runs generated between 7,190-16,804 high quality 2D reads, comprising 11-22.7Mb of sequence, with a mean length of 1,333-1,536bp and an N50 of 1,504-1,801bp (Table 1). The length distribution of these reads (Figure 2) shows a far lower proportion of short sequences than our Trinity assembly of Illumina HiSeq data derived from the same tissue samples, and also improves upon the EST cluster lengths of Casewell et al. [26], derived from pooled venom gland samples from 10 individuals. LAST alignment [28] of the ‘pass’ reads against a Trinity assembly of Illumina HiSeq data (Table 2) suggests a raw error rate in the region of 12% and the majority of errors are insertions or deletions (Table 3) [25, 29]. Based on comparisons of multiple reads from the same transcript, these errors do not appear to be systematic. Since measured current is interpreted by the basecalling software Metrichor as 5mers we also investigated the percentage change in 5mer representation between our MinION data compared to raw and assembled Illumina data for the same samples. Although crude, this analysis reveals under-representation of homopolymer 5mers (Figure 3) [29, 30]. Interestingly, this pattern was not seen when we compared the MinION data to EST sequences derived from Sanger sequencing, nor was there any obvious correlation between the results obtained from Eco6 and Eco8, suggesting that the small size of this dataset (1,070 reads) is complicating these analyses.

**Figure 2.**
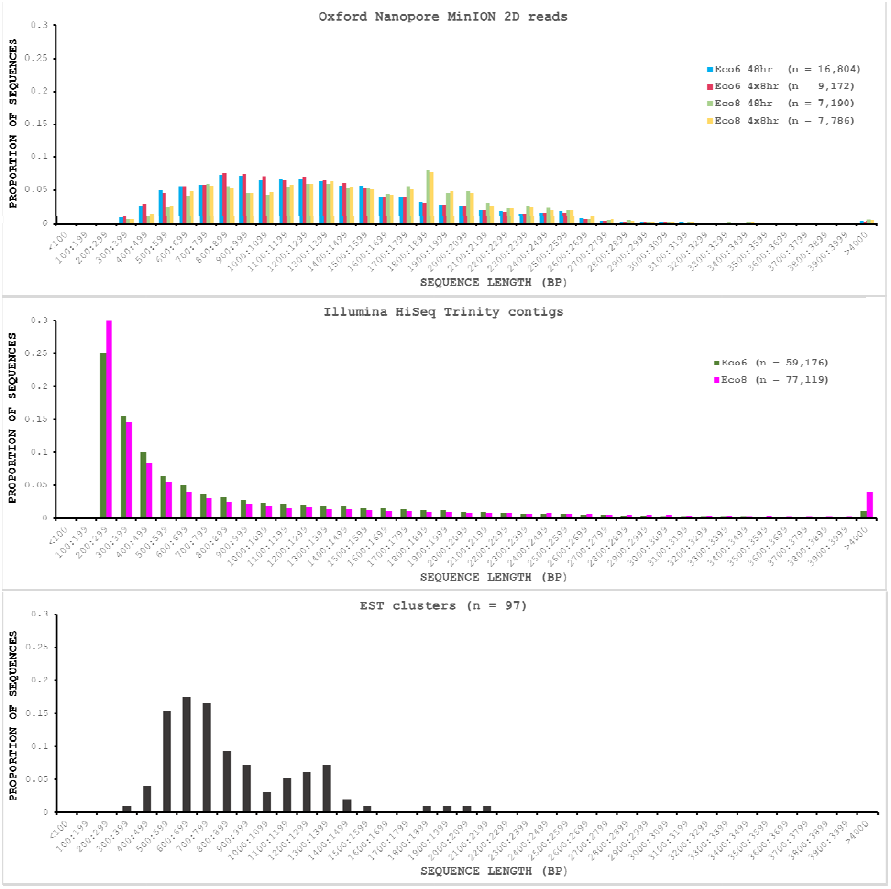
Length distributions of painted saw-scaled viper venom gland sequence data derived from multiple approaches. The Oxford Nanopore MinION data is based only on high quality reads from the Metrichor ‘pass’ folder and is derived from two individuals (Eco6 and Eco8). Both the standard 48 hour sequencing protocol (which performs a re-mux after 24 hours) and a modified protocol with four re-mux steps at 8 hour intervals were used. Illumina HiSeq data derived from the same venom gland tissue samples was assembled using Trinity (version trinityrnaseq_r2012-04-27) and the total number of contigs is indicated for each sample. EST data are from Casewell et al. [26], based on 1,070 Sanger reads, grouped into 97 clusters.

**Figure 3.**
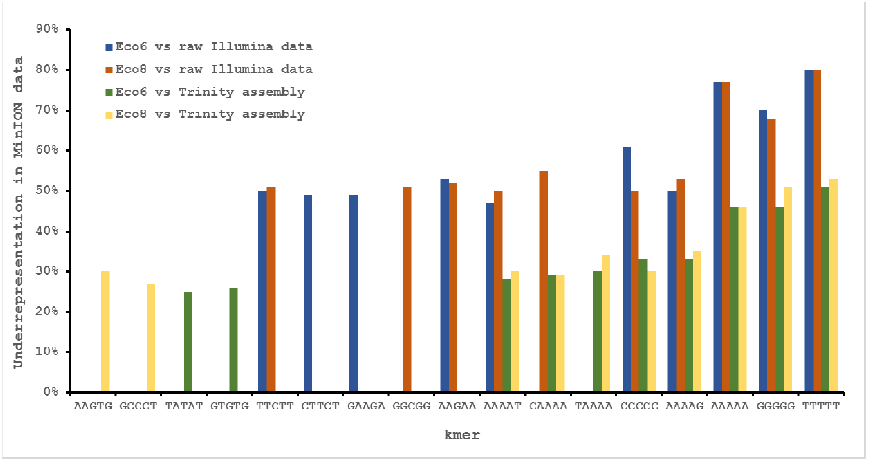
Under-represented kmers in raw Oxford Nanopore MinION data (with pooled runs for each individual) compared to raw and assembled (Trinity version trinityrnaseq_r2012-04-27) Illumina data from the same tissue samples. The ten most under-represented 5mers for each comparison are shown, with homopolymer 5mers particularly under-represented.

**Table 2.** Sequence and assembly statistics for painted saw-scaled viper (*Echis coloratus*) venom gland RNA-Seq and expressed sequence tag (EST) data. Statistics are provided for two *de novo* RNA-Seq assemblers (Trinity and SOAPdenovo-trans [36]) and one genome-guided assembly method (the Tuxedo suite [37]) for which we used a low coverage (∼30x) draft *E. coloratus* genome assembly. EST statistics are based on data from Casewell et al. [26].

|  | Illumina HiSeq<br>Trinity assembly |  | Illumina HiSeq<br>SOAPdenovo-Trans |  | Illumina HiSeq<br>Tuxedo<br>(genome-guided) |  | ESTs |
| --- | --- | --- | --- | --- | --- | --- | --- |
|  | Eco6 | Eco8 | Eco6 | Eco8 | Eco6 | Eco8 |  |
| Number of reads | 52,179,724 |  |  |  |  |  | 1070 |
| Number of bases | 10,513,367,160 |  |  |  |  |  | 676,396 |
| Number of<br>contigs | 59,176 | 77,119 | 136,903 | 169,750 | 33,917 | 48,912 | 97 |
| Max length (bp) | 9,014 | 16,826 | 8,331 | 12,403 | 14,002 | 14,007 | 2,162 |
| N50 (bp) | 1,619 | 2,338 | 1,175 | 2,034 | 1,625 | 1,683 | 652 |

**Table 3.** Correction of Oxford Nanopore MinION sequence derived from the painted saw-scaled viper (*Echis coloratus*) venom gland using proovread and nanocorrect. These approaches reduce the error rate from around 12% to 0–2% and around 4.5% (3% after a second round of correction) respectively. MinION data for the separate runs for the two *E. coloratus* individuals (Eco6 and Eco8) has been pooled.

|  | Uncorrected |  | proovread |  | nanocorrect |  |
| --- | --- | --- | --- | --- | --- | --- |
|  | Eco6 | Eco8 | Eco6 | Eco8 | Eco6 | Eco8 |
| Total reads | 25,976 | 14,976 | 21,751 | 11,066 | 7,357 | 4,762 |
| Total bases (Mb) | 34.9 | 22.7 | 26.1 | 14.6 | 11.4 | 8.1 |
| Length (bp) |  |  |  |  |  |  |
| Max | 12,639 | 8,521 | 5,084 | 5,362 | 4,957 | 4,702 |
| Min | 247 | 248 | 300 | 153 | 19 | 330 |
| N50 | 1,525 | 1,792 | 1,334 | 1,527 | 1,577 | 1,804 |
| Alignment length (bp) | 22,512,857 | 14,029,072 | 24,272,630 | 13,311,248 | 9,948,507 | 6,611,836 |
| Matches | 20,421,908 | 12,647,267 | 24,129,077 | 13,099,487 | 9,623,049 | 6,347,134 |
| Mismatches | 751,390 | 515,464 | 70,120 | 140,336 | 98,888 | 98,961 |
| Insertions | 663,396 | 416,137 | 9,980 | 12,873 | 75,180 | 48,744 |
| Deletions | 1,339,559 | 866,341 | 73,433 | 71,425 | 226,570 | 165,741 |
| Total Errors | 2,754,345<br>**(12.2%)** | 1,797,942<br>**(12.8%)** | 153,533<br>**(0.6%)** | 224,634<br>**(1.7%)** | 400,638<br>**(4.0%)** | 313,446<br>**(4.7%)** |

Hybrid error correction of our MinION reads with higher-quality short read (100bp) Illumina data using proovread [31] reduced the error rate to between 0–2%, with particular reduction in the number of indels relative to mismatches (Table 3). However, for many applications, this type of high coverage short-read data may not be available for error correction, and so we also investigated the feasibility of *de novo* error correction with nanocorrect [30] using only MinION-derived reads for each individual. This approach reduced the error rate to around 4-5% using one round of correction (Table 3), and to around 3% using two rounds, with little to no further improvement seen after subsequent rounds of correction (Supplementary file 1). However, the number of reads post-correction was greatly reduced and many key venom gene families of interest were missing or underrepresented. Finally, we attempted error correction using nanopolish [30], a signal-level consensus algorithm which uses a hidden Markov model to correct assemblies using the original MinION electric current signals, but find that this approach performs poorly compared to both proovread and nanocorrect, giving an error rate of around 7.5%.

To provide some indication of the quality of our Illumina and corrected MinION “assemblies” we used TransRate [32], which assigns overall and optimised quality scores for *de novo* assemblies. An overall score of 0.22 and an optimised score of 0.35 have been suggested to be better than 50% of *de novo* assemblies from NCBI Transcriptome Shotgun Assembly (TSA) database [32]. Our original Trinity assemblies exceed these numbers, as does the Eco8 proovread-corrected dataset (Table 4). The Eco6 proovread-corrected data has an optimised score of 0.42, but an overall score of only 0.13. Whilst it seems likely that the proovread-corrected MinION data quality is similar in quality to those derived from Illumina data, the utility of TransRate for the assessment of corrected MinION “assemblies” will require the analysis of a larger number of datasets, and we include these statistics here mainly for completeness. We next investigated putative protein coding sequences using TransDecoder (version 2.0.1) [33], specifying that any potential open reading frame (ORF) must code for a protein at least 100 amino acids long. The longest putative ORFs were compared to the Swissprot protein database (downloaded on 29/07/2015 from www.uniprot.org) and all ORFs with homology to known proteins retained (Table 4). The corrected MinION data had a higher proportion of predicted mRNAs encoding a ≥100 amino acid protein (Figure 4) and, given the higher values for the proovread-corrected data, and the fact that it contains a greater proportion of key venom gene families, we therefore focussed on this dataset for a more detailed analysis of candidate venom toxin encoding genes in *E. coloratus*.

**Figure 4.**
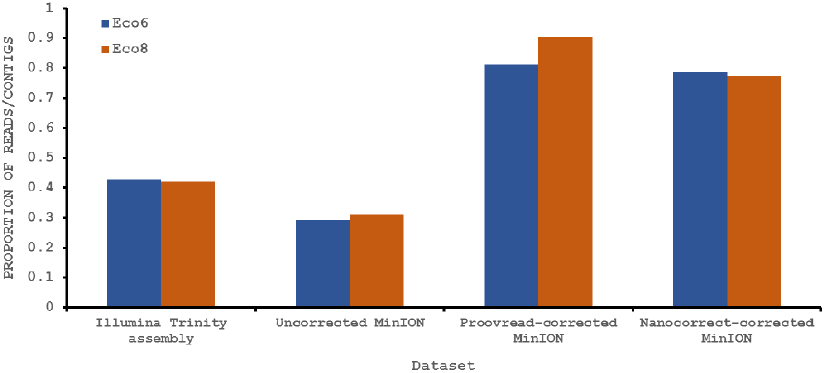
Proportion of Illumina contigs and Oxford Nanopore MinION reads with a predicted mRNA encoding an open reading frame of at least 100 amino acids that has homology to a known protein in the Swissprot protein database. Both hybrid and *de novo* correction greatly increases the proportion of MinION-derived reads with ≥100 amino acid ORF.

**Table 4.** Predicted mRNA sequences and open reading frames (ORFs) as determined by TransDecoder [33] and quality scores as determined by TransRate [32]. MinION data for the separate runs for the two *E. coloratus* individuals (Eco6 and Eco8) has been pooled.

|  | Illumina Trinity assembly |  | Uncorrected Nanopore |  | Proovread-corrected Nanopore |  | Nanocorrect-corrected Nanopore |  |
| --- | --- | --- | --- | --- | --- | --- | --- | --- |
|  | Eco6 | Eco8 | Eco6 | Eco8 | Eco6 | Eco8 | Eco6 | Eco8 |
| Total reads/contigs | 59,176 | 77,119 | 25,976 | 14,976 | 21,751 | 11,066 | 7,357 | 4,762 |
| Read/contig N50 (bp) | 1,492 | 2,142 | 1,525 | 1,792 | 1,334 | 1,527 | 1,577 | 1,804 |
| Predicted mRNAs | 25,395 | 32,424 | 7,587 | 4,628 | 17,685 | 9,985 | 5,779 | 3,679 |
| mRNA N50 (bp) | 2,235 | 3,311 | 1,738 | 1,899 | 1,443 | 1,708 | 1,731 | 1,864 |
| Full length ORFs | 7,985 | 14,867 | 5,339 | 3,461 | 6,616 | 4,564 | 4,409 | 2,887 |
| ORF N50 (aa) | 1,044 | 1,506 | 372 | 375 | 819 | 777 | 405 | 390 |
| TransRate score | 0.25 | 0.35 | 0.06 | 0.07 | 0.13 | 0.32 | 0.05 | 0.09 |
| Optimal score | 0.47 | 0.47 | 0.21 | 0.20 | 0.42 | 0.41 | 0.16 | 0.19 |

We have previously suggested that the venom of *E. coloratus* comprises products from 34 different genes, in 8 gene families [14]. However, in order to gain a better appreciation of the utility of the MinION for characterising venom gland transcriptomes, we have expanded our analyses beyond only these genes to other members of the same gene families which we previously ruled out as contributing to venom toxicity based on low expression levels and/or a wider tissue expression pattern (Figure 5). Our Trinity (version trinityrnaseq_r2012-04-27) assembly of Illumina HiSeq data was able to reconstruct 13/40 full length sequences (which we define as a full open reading frame and at least some 5′ and 3′ untranslated region (UTR) sequence). This number is slightly misleading however, as seven of the *c-type lectin (ctl*) genes have identical 294bp 5′ UTRs and have therefore likely been misassembled, probably as a result of very high similarity in the region encoding the signal peptide. The Sanger-based EST clusters reconstruct 15 of the 29 detected genes (Figure 5). Interestingly, despite its reputation for producing chimeric transcripts [17], we find little evidence of this in our Trinity dataset and in fact encounter such issues only in the EST dataset, where *vegf-f, serine protease b* and *c* and *c-type lectin c* appear to be comprised of concatenated reads. The Illumina-corrected MinION reads for the Eco6 sample provided full coding sequences for 29 of 33 genes detected. Sequence identity between the corrected reads and the Trinity reference was typically 99-100% across the aligned region, and this was often higher than that of the EST clusters, where sequence quality deteriorated towards the ends. We were also able to identify putative splice variants using the MinION data that had not been recovered by either of the other two approaches. Although we did not detect all target transcripts, this was not unexpected for a variety of reasons. Firstly, the Illumina Trinity assembly reference dataset was assembled from several individuals at different time points during venom synthesis following milking and so certain genes may not be expressed in the samples used for our MinION experiments, and secondly, our analysis of the MinION dataset is based on only 40, 952 high-quality reads, whereas the Illumina data for the two samples comprised 52,179,724 paired-end reads (10,513,367,160bp). Investigation of the effect of sequencing depth on the characterisation of snake venom gland transcriptomes using sub-assemblies of existing data (Supplementary file 2) suggests that assemblies based on around 8 million 100bp paired-end reads are able to return BLAST matches to all candidate genes. It is therefore truly exceptional that our much smaller amount of MinION data is able to provide not just matches, but full coding sequences for such a large number of venom genes in our study species. Although developed primarily to boost sequence production at the late stages of flow-cell use, we find that the modified 4x8hr run scripts produce a much smoother data acquisition profile (Figure 6) and it seems likely that further refinements in this area will greatly improve data generation. The largest contributor to total sequence output however seems to be the number of available pores on each flowcell (Table 1, Figure 6) and greater consistency in this area, together with planned future increases to the number of pores per flowcell and the speed at which DNA traverses the pore will greatly increase the amount of data generated per flowcell. As an example of the speed at which the MinION and its associated technology and reagents are developing, we used the latest versions of Metrichor and 2D basecalling workflow (version 2.26.1 and 1.14 respectively) to re-analyse the Okinawa habu (*Protobothrops flavoviridis*) venom gland data that Mikheyev and Tin [25] produced using an amplicon sequencing kit (most likely DEV-MAP001) and R6 flowcells. Despite less than a year separating our and their experiments, the runs that we performed using R7.3 flowcells using the 2D cDNA sequencing protocol with Nanopore Sequencing Kit SQK-MAP005 generated (roughly) 20-45 times as many reads; 50-100Mb more total sequence; 450-1000 times as much high-quality data and 500-1000 times as much high-quality sequence. These figures clearly demonstrate the rapid pace of development of the Oxford Nanopore MinION.

**Figure 5.**
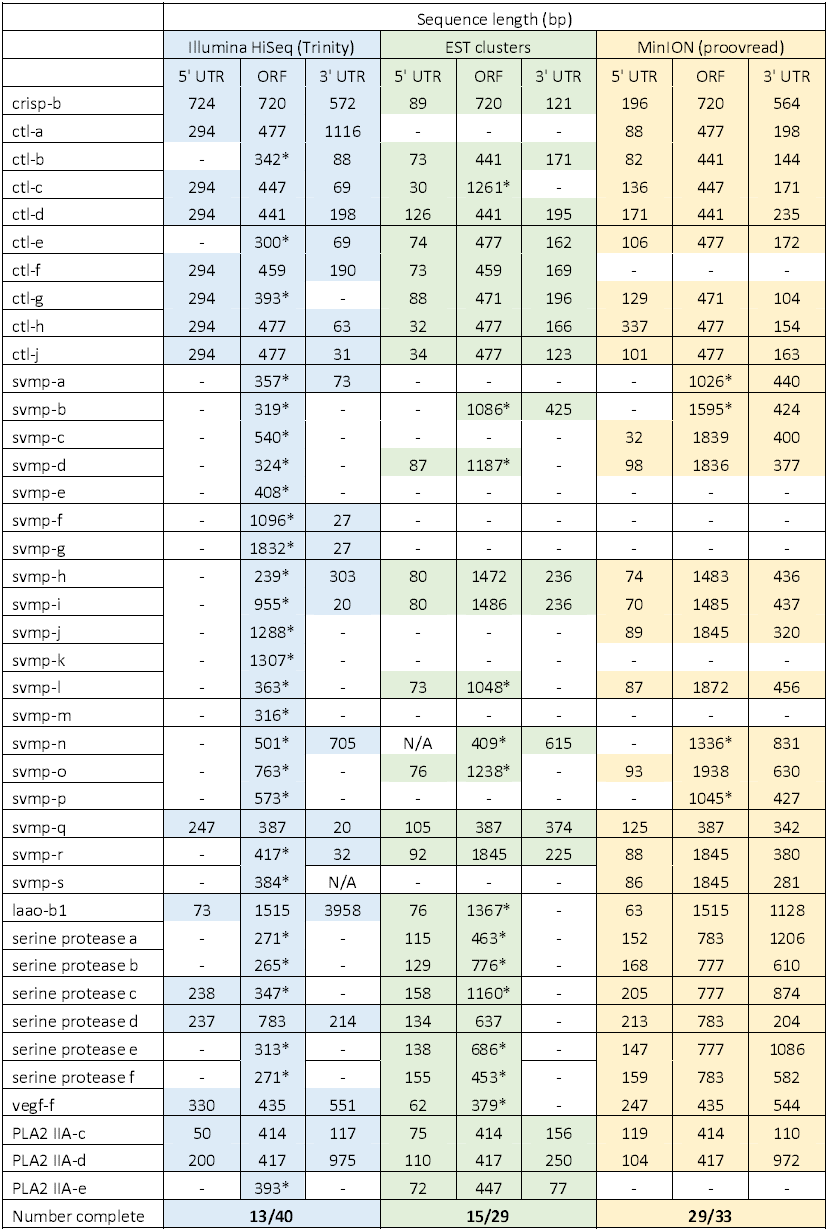
Comparisons of different sequencing approaches for the characterisation of transcripts encoding venom toxins in the painted saw-scaled viper (*Echis coloratus*) venom gland. The reference set of 40 candidate venom genes is derived from a Trinity (version trinityrnaseq_r2012-04-27) assembly of Illumina HiSeq data, where 13 transcripts contain the full open reading frame (ORF), although it is likely that the true number is lower, as the identical 5′ UTR length of c-type lectin (ctl) transcripts suggests misassembly. A set of EST clusters derived from 1,070 Sanger sequences from a pool of 10 individuals [26] detects 29 of these transcripts, 15 of which contain the full ORF. Data generated using the Oxford Nanopore MinION, corrected using proovread, is able to detect 33 candidates, of which 29 contain the full ORF. Incomplete ORFs are indicated with an *.

**Figure 6.**
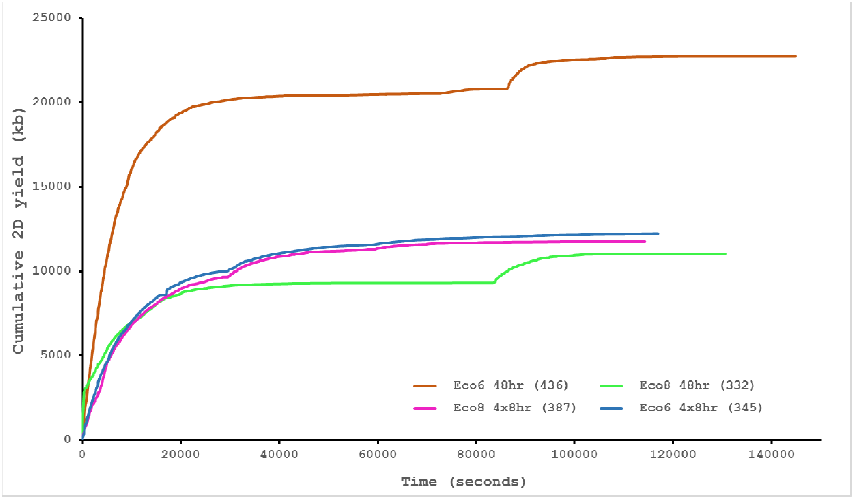
Data acquisition of 2D reads during four runs of the Oxford Nanopore MinION, using cDNA derived from two individuals (Eco6 and Eco8). Both the standard 48 hour protocol (including a remux after 24 hours) and a set of modified run scripts that perform four re-muxes at 8 hour intervals were used. This latter approach yields a much smoother data acquisition profile. The number of pores available at initial QC for each flowcell is given in brackets in the legend.

## Conclusions

Until relatively recently it seemed as though DNA sequencing was coming to be dominated by a single company, and a single platform (or at the very least, a closely related family of platforms), with a particular focus on generating an ever-increasing number of human genome sequences. Indeed, the Illumina HiSeq X Ten system has been engineered to *only* be able to sequence human genomes and the required $10 million outlay restricts the number of potential purchasers significantly. Benchtop systems such as the Illumina MiSeq are more affordable and are becoming increasingly common at the research group or institutional level, although they still require a not-insignificant initial outlay and ongoing maintenance and update programs. Against this background, the Oxford Nanopore MinION has the potential to be a truly disruptive technology, offering long reads (in theory limitless, but in practise determined by the size of DNA fragments provided by the user), low and flexible pricing (including a “Zero Hour Flowcell” plan, where users can pay lower amounts for a defined number of hours of sequencing) and portability. This latter is particularly important for field-based species identification, or for rapid response to disease outbreaks [34]. Planned or ongoing updates to the MinION, such as the release of the MinION MkI, new flowcells with increased numbers of pores, “fastmode” sequencing to increase output and automated sample preparation techniques will go some way to enabling the MinION to meet its full potential, but we predict that the greatest advances will come from improvements to the basecalling algorithms. However, hybrid approaches combining MinION data with shorter, more accurate Illumina reads are clearly already effective and can produce a fully circularised bacterial genome for around £500 [35], and *de novo* error-correction approaches have been shown to be possible in at least some cases [30]. For our purposes, a hybrid approach to error correction provided full coding sequences for a large number of venom toxin encoding genes, and was superior to both Illumina-only approaches and Sanger-based ESTs. We therefore suggest that, in the absence of reference genomes, such hybrid approaches will become the default method for the characterisation of transcriptomes from a wide range of species.

## Methods

### mRNA extraction and double-stranded cDNA synthesis

Total RNA was extracted from the venom glands of two *Echis coloratus* (snap-frozen after removal and stored at −80°C [14, 15]) using TriReagent (Sigma T9424) and mRNA purified using the polyA Spin mRNA Isolation Kit (New England BioLabs S1560). mRNA was quantified using a Qubit fluorometer (Qubit RNA HS Assay Kit Q32852) and reverse transcription carried out using 120ng (Eco6) or 240ng of mRNA (Eco8). Primer annealing was performed at 65°C for 5 minutes in a 13μl reaction comprising the required amount of mRNA, 2μl of 1μl Oligo d(T)23 VN primer (New England BioLabs S1327S), 1μl of 10mM dNTPs and the appropriate volume of Rnase-free water. The reaction was then snap-cooled on a pre-chilled freezer block. 4μl of 5× First Strand buffer and 2μl 100 mM DTT (part of Life Technologies 18064-014) were then added to the primer/mRNA mix, which was briefly vortexed, spun down in a microcentrifuge and incubated at 42°C for 2 minutes. Finally, 1 μl of 200 U/μl SuperScript II Reverse Transcriptase (Life Technologies 18064-014) was added to each tube and reverse transcription carried out at 50°C for 50 minutes, with a subsequent 15 minute incubation at 70°C for enzyme denaturation. Second strand synthesis was performed with the NEBNext mRNA Second Strand Synthesis Module (New England BioLabs E6111), using 45μl of nuclease-free water, 10μl of NEBNext Second Strand Synthesis Reaction Buffer and 5μl of NEBNext Second Strand Synthesis Enzyme Mix, with incubation at 16°C for 1 hour. Double-stranded cDNA (ds cDNA) was purified using a 1.8× volume of Agencourt AMPure XP beads (Beckman Coulter A63880), with a 5 minute binding step (with gentle shaking), two washes in 200μl 70% ethanol and elution in 51μl nuclease-free water.

### End-repair and dA-tailing

End-repair was performed using the NEBNext End Repair Module (New England BioLabs E6050) with 6μl of 10× end-repair buffer and 3μl of end-repair enzyme mix added to each of the 51¼l ds cDNA samples, followed by incubation at room temperature for 25 minutes and clean-up using a 1.8× volume of Agencourt beads (as above), with elution in 25μl nuclease-free water. Next, the end repaired ds cDNA was dA-tailed with the NEBNext dA-Tailing Module (New England BioLabs E6053), using 3μl of 10× NEBNext dA-Tailing Reaction Buffer and 2μl of A-tailing enzyme (Klenow Fragment (3′→ 5′ exo-)) and incubation at 37°C for 30 minutes, followed by clean-up with 1.8× Agencourt beads (as above) and elution in 15μl of nuclease-free water.

### PCR adapter ligation and amplification

Prior to amplification, adapters were ligated to the end-repaired, dA-tailed ds cDNA using 5μl of the Oxford Nanopore SQK-MAP005 PCR adapters (a double-stranded oligonucleotide supplied by Oxford Nanopore, formed by heating a solution containing each oligo (Short_Y_top_LI32 5′-GGTTGTTTCTGTTGGTGCTGATATTGCGGCGTCTGCTTGGGTGTTTAACCT-3′ and Y_bottom_LI33 5′-

GGTTAAACACCCAAGCAGACGCCGAAGATAGAGCGACAGGCAAGTTTTGAGGCGAGC GGTCAA-3′) at 20μM in 50 mM NaCl, 10 mM Tris-HCl pH7.5 to 95°C for 2 minutes, and cooling by 0.1°C every 5 seconds) and 20μl of Blunt/TA Ligase Master Mix (New England BioLabs M0367), with incubation at room temperature for 15 minutes. Adapter-ligated DNA was purified using 0.7x of Agencourt beads (as above) and eluted in 25μl nuclease-free water, followed by amplification using 50μl of LongAmp Taq 2× master mix (New England BioLabs M0287), 2μl of Oxford Nanopore SQK-MAP005 PCR primers (PR2 5′-TTTCTGTTGGTGCTGATATTGC-3′ and 3580F 5′-ACTTGCCTGTCGCTCTATCTTC-3′) and 23μl nuclease-free water. Initial denaturation was 95°C for 3 minutes, followed by 15 cycles of 95°C for 15 seconds, 62°C for 15 seconds and 65°C for 5 minutes, with a final extension at 65°C for 10 minutes. Amplified DNA was purified using 0.7× Agencourt beads (as above) with elution in 80μl of nuclease-free water.

### Sequencing adapter ligation

End-repair of the amplified DNA was carried out using the NEBNext End Repair Module (New England BioLabs E6050), with 10μl of 10× end-repair buffer, 5μl of end-repair enzyme mix and 5μl of nuclease-free water and incubation at room temperature for 20 minutes. End-repaired DNA was purified using 1× volume of Agencourt beads as outlined previously, with elution in 25μl of nuclease-free water. dA-tailing and clean-up was carried out as described above, with elution in 30μl of nuclease-free water. Adapter ligation was performed for 10 minutes at room temperature in Protein LoBind 1.5 ml Eppendorf tubes (Sigma Aldrich Z666505-100EA) using 10μl of each of the Oxford Nanopore SQK-MAP005 adapter and HP adapters and 50μl of Blunt/TA Ligase Master Mix (New England BioLabs M0367). Clean-up was performed using an equal volume of Dynabeads His-Tag Isolation and Pulldown beads (Life Technologies 10103D), which had been washed twice in SQK-MAP005 1× Bead Binding Buffer and resuspended in 100μl of 2× Bead Binding Buffer. The bead/DNA mix was incubated at room temperature for 5 minutes to allow binding, washed twice in 200μl of 1× Bead Binding Buffer, eluted in 25μl of elution buffer and the resulting ‘Pre-sequencing library’ either used immediately or stored at −20°C in 6μl aliquots in LoBind tubes.

### Flowcell preparation and sample loading

A total of four Oxford Nanopore FLO-MAP003 (R7.3) flowcells were used, and these were stored at 4°C from delivery until use. Flowcells were fitted into MIN-MAP001 MinION Sequencing Devices and secured using the provided nylon screws and new heat pads were used for each flowcell. Prior to sample loading, the flowcells were primed using two 10 minute washes of 150μl of 1× SQK-MAP005 Running Buffer with 3.25μl of Fuel Mix. Finally, a 6μl aliquot of the pre-sequencing library was mixed with 75μl of 2× Running Buffer, 66μl of nuclease-free water and 3μl of Fuel Mix then briefly mixed by inversion, microfuged and loaded onto the flowcell.

### Sequencing

Sequencing utilised both the standard 48-hour sequencing protocol and a modified 4x 8-hour protocol (J. Tyson, pers com.), run using the MinKNOW software (version 0.49.2.9). For the 48hr runs, a fresh aliquot of sequencing library was added at around 24 hours. Base-calling from read event data was performed by Metrichor (version 2.26.1) using the 2D basecalling workflow (version 1.14). We also re-analysed the Okinawa habu (*Protobothrops flavoviridis*) venom gland data of Mikheyev and Tin [25] using this Metrichor version and workflow.

### Data analysis

Sequencing statistics were determined and data extracted in .fastq and .fasta format using poretools [38] and poRe [39]. Error correction was carried out using both hybrid and *de novo* correction methods. Hybrid error correction using short-read (2x100bp paired-end reads) sequencing data previously generated on the Illumina HiSeq platform was carried out using a module of proovread [31]. More specifically, we utilised proovread-flex, which is optimised for the uneven sequencing coverage seen in metagenomes and transcriptomes. For *de novo* error correction we utilised nanocorrect [30] (available at https://github.com/jts/nanocorrect) using commands based on the full pipeline script found at https://github.com/jts/nanopore-paper-analysis/blob/master/full-pipeline.make. A single round of correction was carried out for each individual and multiple rounds trialled on Eco6 data only. We also used nanopolish [30], which corrects based on the electrical signal events recorded in the original .fast5 file of the MinION read, using commands found at https://github.com/jts/nanopolish. Sequence accuracy was assessed using BWA-MEM [40] alignments and python scripts found at https://github.com/arq5x/nanopore-scripts following Loman et al [30], assembly quality was determined using TransRate [32] and putative protein-coding open-reading frames predicted using TransDecoder [33]. Corrected reads of interest were identified with BLAST+ (version 2.2.29 [41]) using query sequences from a previously generated reference venom gland transcriptome assembly [14, 15]. Sequences were aligned using CLUSTAL [42] and manually annotated to identify the protein coding ORF and 5′ and 3′ UTRs.

### Data access

Raw MinION venom gland data has been deposited in the European Nucleotide Archive under study number PRJEB10285 (Eco6 48hr run ERR985427; Eco6 4x8hr run ERR986484; Eco8 48hr run ERR985428; Eco8 4x8hr run ERR985429) and previously generated short-read sequencing data for Eco6 and Eco8 venom gland samples [14, 15] can be obtained from the SRA database under the accessions ERS094900 and SRX543069 respectively.

## Acknowledgements

The authors wish to thank Oxford Nanopore for letting us onto the MinION Access Program, and Mick Watson, Thomas Hackl, Nick Loman and Jared Simpson for various help with poRe, poretools, proovread and nanocorrect. We would also like to thank John Tyson for providing access to modified run scripts via the Oxford Nanopore discussion space and Alexander Mikheyev for the *Protobothrops flavoviridis* raw data. We are also grateful to the staff of High Performance Computing (HPC) Wales and Peter Holland for enabling and supporting our access to their systems. This research was partially supported by a Royal Society Research Grant awarded to JFM (grant number RG100514) and JFM has also been generously supported by the Biosciences, Environment and Agriculture Alliance (BEAA) between Bangor University and Aberystwyth University.

## Supplementary file 1 : *de novo* error correction using Nanocorrect

**Supplementary Table S1.** Correction of pooled Eco6 nanopore sequence data using up to five rounds of correction using Nanocorrect [1]. Little improvement is seen after the second round, although the number of sequences is much reduced as a result of this process.

|  | Nanocorrect correction |  |  |  |  |
| --- | --- | --- | --- | --- | --- |
|  | <u>Round 1</u> | <u>Round 2</u> | <u>Round 3</u> | <u>Round 4</u> | <u>Round 5</u> |
| Alignment length (bp) | 9,948,507 | 5,807,383 | 4,245,548 | 3,681,607 | 3,363,671 |
| Matches | 9,623,049 | 5,660,507 | 4,140,907 | 3,591,236 | 3,280,638 |
| Mismatches | 98,888 | 40,682 | 31,202 | 26,821 | 25,011 |
| Insertions | 75,180 | 26,494 | 18,393 | 15,983 | 14,799 |
| Deletions | 226,570 | 106,194 | 73,439 | 63,550 | 58,022 |
| Total Errors | 400,638<br>**(4.0%)** | 173,370<br>**(3.0%)** | 123,034<br>**(2.9%)** | 106,354<br>**(2.9%)** | 97,832<br>**(2.9%)** |

## Supplementary file 2 - Sub-assemblies of the *Echis coloratus* venom gland transcriptome

In an attempt to determine the minimum required amount of sequencing to fully sequence and assemble the venom gland transcriptome of *Echis coloratus*, sub-sets of RNA-seq reads were extracted and assembled (Table S2). Paired venom gland reads were first interleaved using the `shuffleSequences.pl` perl script (part of the Velvet *de novo* assembly program [1]) so that each read pair was maintained during sub-sampling. Using the commands head and tail, 3 sub-sets (designated as “head”, “middle” and “tail”) of either 2, 4, 8 or 10 million reads were taken from an RNA-seq dataset containing 44,678,609 paired-end reads. These data were assembled using Trinity [2,3], with parameters set to run as a single-end read dataset (as there is only one .fastq input file), but with the added command-line parameter `- -run_as_paired` to indicate that the data contains paired-end data.

**Supplementary Table S2.** Assembly metrics for sub-assemblies of the venom gland transcriptome of *Echis coloratus*.

| Sub-sample | Sample size (million reads) | Total number of contigs | Number of contigs $\geq 300$nt | Total length (nt) | Max. contig size (nt) | Contig N50 (nt) |
| --- | --- | --- | --- | --- | --- | --- |
| H<br>E<br>A<br>D | 2 | 24,585 | 14,744 | 10,302,850 | 7,474 | 808 |
|  | 4 | 34,990 | 22,184 | 17,605,771 | 7,860 | 1,023 |
|  | 8 | 45,207 | 30,121 | 27,542,537 | 11,824 | 1,293 |
|  | 10 | 48,349 | 32,660 | 31,623,176 | 11,824 | 1,420 |
| M<br>I<br>D<br>D<br>L<br>E | 2 | 23,915 | 14,229 | 10,036,594 | 7,840 | 837 |
|  | 4 | 34,383 | 21,736 | 17,282,856 | 8,970 | 1,027 |
|  | 8 | 44,759 | 29,946 | 27,116,697 | 11,738 | 1,279 |
|  | 10 | 47,832 | 32,451 | 30,985,872 | 11,752 | 1,387 |
| T<br>A<br>I<br>L | 2 | 24,170 | 14,513 | 10,059,952 | 8,547 | 810 |
|  | 4 | 34,735 | 21,994 | 17,315,514 | 8,165 | 1,004 |
|  | 8 | 44,956 | 29,988 | 27,283,356 | 11,803 | 1,284 |
|  | 10 | 48,116 | 32,535 | 31,022,314 | 11,805 | 1,382 |

Local blast surveys were then carried out using BLAST+ version 2.2.27 [4] to identify previously characterised putative toxin genes in *E. coloratus*. The majority of transcripts encoding putative toxin genes appear to be present in venom gland transcriptome assemblies generated from only 2 million paired-end reads (here presence is defined as the transcript being found in all three (Head/Middle/Tail) sub-assemblies) (Table S3).

**Supplementary Table S3.** Presence/absence of putative toxin transcripts in sub-assemblies of the venom gland transcriptome of *Echis coloratus*. Detected transcripts are shaded, transcripts not found are shaded grey. H, head; M, middle; T, tail.

| Gene | Sub-sample size and position |  |  |  |  |  |  |  |  |  |  |  |
| --- | --- | --- | --- | --- | --- | --- | --- | --- | --- | --- | --- | --- |
|  | 2 million reads |  |  | 4 million reads |  |  | 8 million reads |  |  | 10 million reads |  |  |
|  | H | M | T | H | M | T | H | M | T | H | M | T |
| 3ftx-a | + | - | + | + | + | + | + | + | + | + | + | + |
| 3ftx-b | + | + | + | + | + | + | + | + | + | + | + | + |
| ache - transcript 1 | - | + | - | + | + | + | + | + | + | + | + | + |
| complement c3 | + | + | + | + | + | + | + | + | + | + | + | + |
| crisp-b | + | + | + | + | + | + | + | + | + | + | + | + |
| crotamine-like | + | + | + | + | + | + | + | + | + | + | + | + |
| c-type lectins a-k | + | + | + | + | + | + | + | + | + | + | + | + |
| cystatin e/m | + | + | + | + | + | + | + | + | + | + | + | + |
| dpp 3 | + | + | + | + | + | + | + | + | + | + | + | + |
| dpp 4 | + | + | + | + | + | + | + | + | + | + | + | + |
| esp-e1 | + | + | + | + | + | + | + | + | + | + | + | + |
| ficolin | + | + | + | + | + | + | + | + | + | + | + | + |
| kallikrein | + | + | + | + | + | + | + | + | + | + | + | + |
| kunitz 1 | + | + | + | + | + | + | + | + | + | + | + | + |
| kunitz 2 | + | + | + | + | + | + | + | + | + | + | + | + |
| laao-a | + | + | + | + | + | + | + | + | + | + | + | + |
| laao-b1 | + | + | + | + | + | + | + | + | + | + | + | + |
| laao-b2 | + | + | + | + | + | + | + | + | + | + | + | + |
| lipa-a | + | + | + | + | + | + | + | + | + | + | + | + |
| lipa-b | - | - | - | + | + | + | + | + | + | + | + | + |
| ngf | + | + | + | + | + | + | + | + | + | + | + | + |
| PLA<sub>2</sub> IIA-c | + | + | + | + | + | + | + | + | + | + | + | + |
| PLA<sub>2</sub> IIA-d | + | + | + | + | + | + | + | + | + | + | + | + |
| PLA<sub>2</sub> IIA-e | + | + | + | + | + | + | + | + | + | + | + | + |
| PLA<sub>2</sub> IIE | - | - | - | - | + | - | - | + | + | - | - | + |
| plb | + | + | + | + | + | + | + | + | + | + | + | + |
| renin | + | + | + | + | + | + | + | + | + | + | + | + |
| serine protease a-f | + | + | + | + | + | + | + | + | + | + | + | + |
| svmp-a | + | + | + | + | + | + | + | + | + | + | + | + |
| svmp-b | + | + | + | + | + | + | + | + | + | + | + | + |
| svmp-c | - | - | + | - | + | + | + | + | + | + | + | + |
| svmp-d | - | - | - | - | + | + | + | + | + | + | + | + |
| svmp-e | + | + | + | + | + | + | + | + | + | + | + | + |
| svmp-f | + | + | + | + | + | + | + | + | + | + | + | + |
| svmp-g | + | + | + | + | + | + | + | + | + | + | + | + |
| svmp-i | + | + | + | + | + | + | + | + | + | + | + | + |
| svmp-j | + | + | + | + | + | + | + | + | + | + | + | + |
| svmp-k | + | + | + | + | + | + | + | + | + | + | + | + |
| svmp-m | + | - | + | + | + | + | + | + | + | + | + | + |
| svmp-n | + | + | + | + | + | + | + | + | + | + | + | + |
| svmp-p | + | - | + | + | + | + | + | + | + | + | + | + |
| svmp-q | + | + | + | - | - | + | + | + | + | + | + | + |
| svmp-t | + | + | + | + | + | + | + | + | + | + | + | + |
| vegf-a | - | - | - | + | - | + | + | + | + | + | + | + |
| vegf-c | - | - | - | - | - | - | + | - | + | + | + | + |
| vegf-f | + | + | + | + | + | + | + | + | + | + | + | + |
| waprin | + | - | + | + | + | + | + | + | + | + | + | + |

As the number of reads used for assembly increases the mean length of the amino acid sequence encoded by the assembled transcript also increases, although there is only a 36 amino acid increase between 2 million and 10 million reads (Figure S1).

**Supplementary figure S1.**
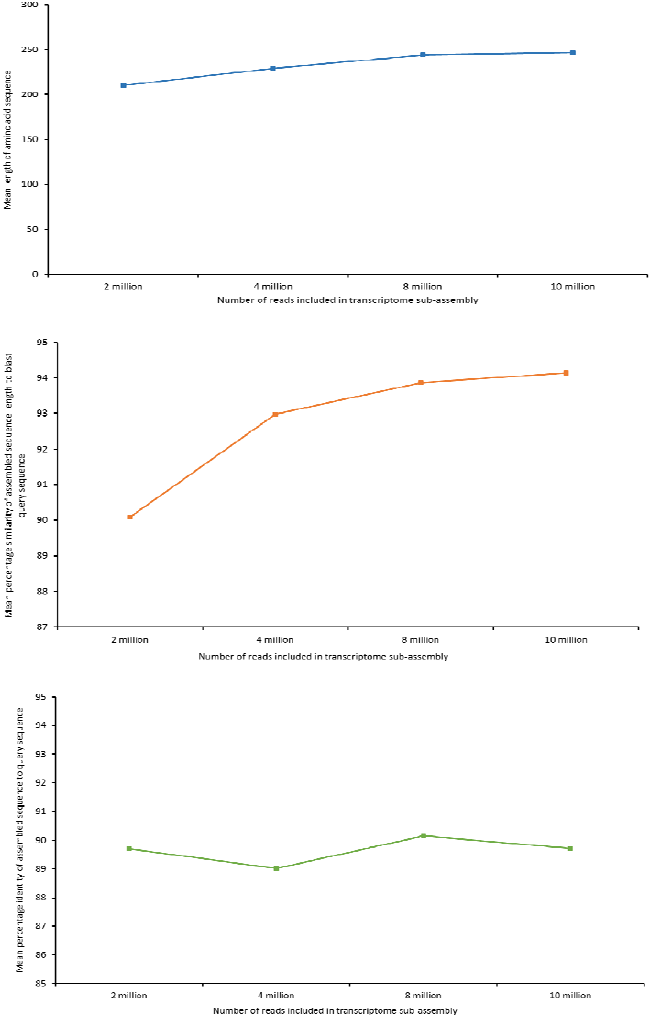
Analysis of sequence assembly quality based on local blast surveys using previously characterised amino acid sequences from *Echis coloratus* venom gland. Top - mean length of amino acid sequence matches in sub-assemblies, Middle - mean percentage length of query sequence covered by assembled sequence. Bottom - mean percentage similarity of assembled sequence to query sequence in sub-assemblies.

However, the number of contigs 300≥bp roughly doubles (Table S1), meaning considerably fewer contigs which are likely to be unplaced paired reads are present in the transcriptome assembly. To gain insight into how this increase in length relates to the quality of the assembled toxin transcript sequences, the percentage of the query sequence covered by the newly assembled sequence was calculated. Again there is only a minor improvement of 4% following an increase from 2 million reads to 10 million (Figure S1). The mean percentage similarity between assembled sequence and query sequence appears to be more variable across the sub-assemblies, with no apparent consistent improvement as the number of reads increases (Figure S1). As the query sequences used for local BLAST searches were obtained from an assembly of multiple *E. coloratus* venom gland datasets in order to represent an overabundance of sequencing, and the sub-assemblies were assembled from a different set of venom gland reads, it should be expected that not all blast alignments will have a 100% match between query and subject due to variation between individuals. However, a lower % identity would indicate that either sequencing errors were incorporated into the assembly or there has been a misassembly, both likely due to a reduced depth of sequencing coverage.

## Conclusion

Around 8 million reads appears to be sufficient sequencing depth to capture all putative toxin-encoding transcripts to a suitable assembly quality. The Illumina HiSeq2500 sequencing platform can currently produce 300–400 million 100nt paired-end reads in “high output” mode, or 200–300 million 150nt paired-end reads in “rapid run” mode. With this in mind, and 8 million paired-end reads assumed to be the minimum sequencing depth required to fully capture all putative toxin transcripts, it is possible to sequence ∼40 venom gland libraries on one sequencing lane of the Illumina HiSeq2500 (in “high output” mode).

